# Using Network Analysis to Localize the Epileptogenic Zone from Invasive EEG Recordings in Intractable Focal Epilepsy

**DOI:** 10.1101/247387

**Authors:** Adam Li, Bhaskar Chennuri, Sandya Subramanian, Robert Yaffe, Steve Gliske, William Stacey, Robert Norton, Austin Jordan, Kareem A. Zaghloul, Sara K. Inati, Shubhi Agrawal, Jennifer J. Haagensen, Jennifer Hopp, Chalita Atallah, Emily Johnson, Nathan Crone, William S. Anderson, Zach Fitzgerald, Juan Bulacio, John T. Gale, Sridevi V. Sarma, Jorge Gonzalez-Martinez

**Author notes:** First two authors contributed equally (Adam Li, Bhaskar Chennuri). Corresponding author: Sridevi V. Sarma.

## Abstract

Treatment of medically intractable focal epilepsy (MIFE) by surgical resection of the epileptogenic zone (EZ) is often effective provided the EZ can be reliably identified. Even with the use of invasive recordings, the clinical differentiation between the EZ and normal brain areas can be quite challenging, mainly in patients without MRI detectable lesions. Consequently, despite relatively large brain regions being removed, surgical success rates barely reach 60-65%. Such variable and unfavorable outcomes associated with high morbidity rates are often caused by imprecise and/or inaccurate EZ localization. We developed a localization algorithm that uses network-based data analytics to process invasive EEG recordings. This network algorithm analyzes the centrality signatures of every contact electrode within the recording network and characterizes contacts into susceptible EZ based on the centrality trends over time. The algorithm was tested in a retrospective study that included 42 patients from four epilepsy centers. Our algorithm had higher agreement with EZ regions identified by clinicians for patients with successful surgical outcomes and less agreement for patients with failed outcomes. These findings suggest that network analytics and a network systems perspective of epilepsy may be useful in assisting clinicians in more accurately localizing the EZ.

**AUTHOR SUMMARY:** Epilepsy is a disease that results in abnormal firing patterns in parts of the brain that comprise the epileptogenic network, known as the epileptogenic zone (EZ). Current methods to localize the EZ for surgical treatment often requires observations of hundreds of thousands of EEG data points measured from many electrodes implanted in a patient’s brain. In this paper, we used network science to show that EZ regions may exhibit specific network signatures before, during and after seizure events. Our algorithm computes the likelihood of each electrode being in the EZ and tends to agree more with clinicians during successful resections and less during failed surgeries. These results suggest that a networked analysis approach to EZ localization may be valuable in a clinical setting.

## INTRODUCTION

Epilepsy is one of the most common brain disorders, characterized by chronically recurrent seizures resulting from excessive electrical discharges from groups of neurons (8). Epilepsy affects over 50 million people worldwide and over 30% of all individuals with epilepsy have intractable seizures, which cannot completely be controlled by medical therapy (3; 4; 35). That is, seizures continue to occur despite treatment with a maximally tolerated dose of at least two anti-epilepsy drugs (AEDs). The direct cost of assessing and treating patients with medically intractable focal epilepsy (MIFE) ranges from $3-4 billion annually ($16 billion in direct and indirect costs) in the US (41). 80% of these costs are incurred by patients whose seizures are not adequately controlled by AEDs (2). The burden of MIFE, however, is much greater than heavy financial costs. MIFE is a debilitating illness where individuals lose their independence, causing profound behavioral, psychological, social, financial and legal issues (14; 16; 17; 23; 49). Cognitive performance may be impaired by MIFE as well as by side effects of AED therapy (14; 16; 17; 23; 49).

Despite the heavy sequelae from MIFE, there is a potentially curative procedure - surgical resection of the epileptogenic zone (EZ), which can be defined as the minimal area of brain tissue responsible for generating the recurrent seizure activity (36). However, to be effective, this procedure depends on correct anatomical identification of the EZ, which is often poorly defined. A comprehensive pre-surgical evaluation is necessary to better delineate the EZ as well as to identify the risk of neurologic morbidity such as motor, visual, or speech impairment. Various non-invasive methods are currently applied in the attempt of defining the EZ, the eloquent cortical and subcortical areas and, consequently, the optimal resective surgical strategy. Non-invasive techniques include scalp EEG and video-EEG monitoring, neuropsychological tests, speech-language studies, and brain imaging (MRI, PET, Ictal SPECT). Of these methods, the highest predictor of surgical success is identification of a single visible MRI lesion (9; 26; 27; 40; 50; 54).

Localization and surgical success in seizure control are even more challenging in patients with non-lesional MRI. When the non-invasive methods of localization fail to identify the EZ, an invasive monitoring evaluation may be indicated, involving the implantation of subdural grid electrodes (SDE) through open craniotomies or stereo-electroencephalography (SEEG) (42; 44; 59). The process of identifying the EZ then involves visually inspecting tens to hundreds of invasive EEG signals without much assistance from computational tools. Epileptologists currently study the onset of seizure events that occur over several days. Early presence of beta-band activity (beta buzz) or bursts of high frequency oscillations (HFOs) in the 100-300 Hz range, which typically occur milliseconds before the clinical onset of seizures are localizing of the seizure onset (15). Channels where seizure onset features first appear are commonly defined as the seizure onset zone (SOZ), the current best estimate of the unknown EZ. This is based on the assumption that the epileptic cortex generates epileptiform activity, which then entrains other regions into a clinical seizure (15). Electrodecremental responses (loss of rhythmic activity) are also often observed. In general, epileptologists look at a variety of signatures to make their decision (15). Despite all of these possible EEG signatures, determination of the EZ may remain unclear for non-lesional patients (20; 29; 43; 60). See Fig. 1 for a schematic of a current clinical process of localizing the EZ.

**Figure 1.**
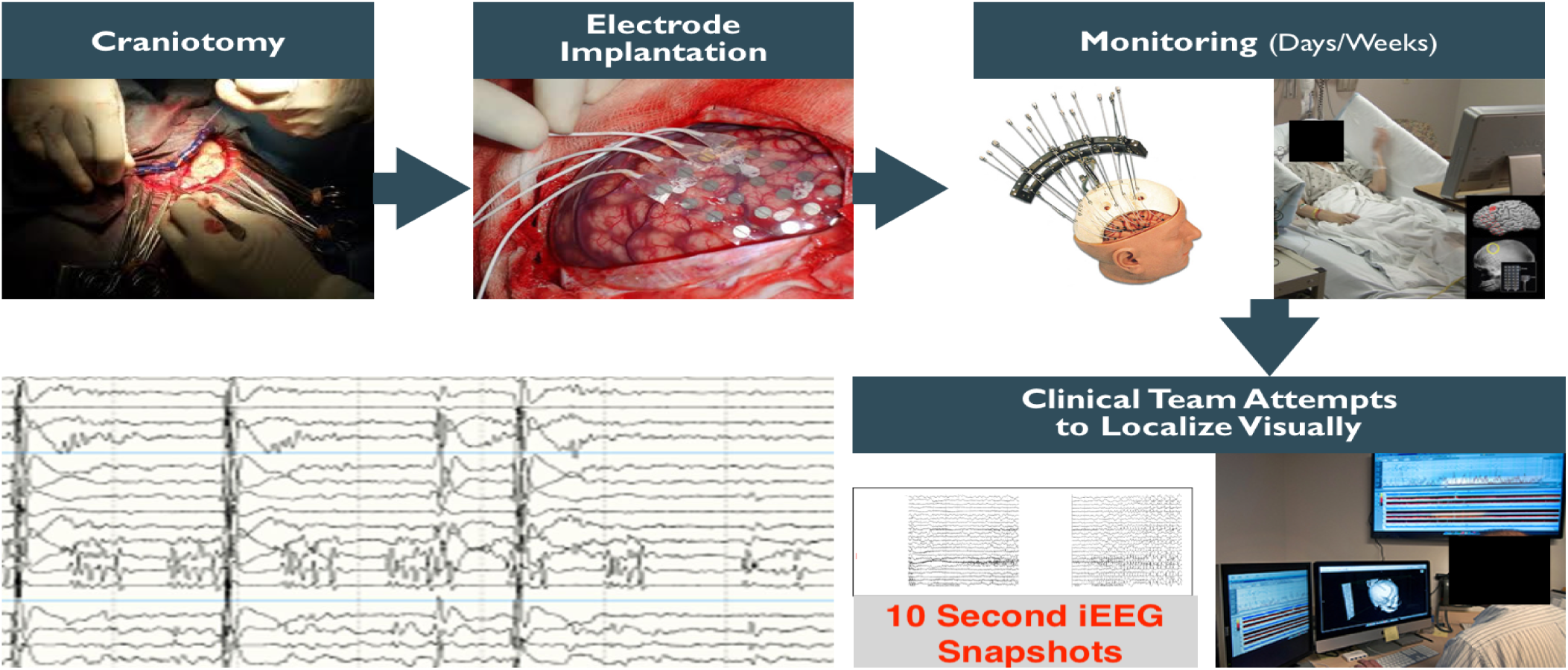
Clinical process for implantation of SDE and seizure onset localization. Clinicians expose the brain through a craniotomy, then implant electrodes on the cortical surface of the brain, monitor patient electrocorticography (ECoG) for days/weeks and then attempt to localize the EZ visually. Clinical teams look at recorded data on computers and annotate signals from certain electrodes and time periods.

Network analysis of intracranial EEG data has been heavily used to study brain activity (1; 7; 10; 13). Networked-based analysis assumes that signals from different EEG channels are samples of activity from brain regions that are structurally and/or functionally connected and therefore dependent (30; 46; 63). Several important prior studies have looked at network dynamics in epileptic cortex during seizure events. Some works investigate correlation structure over seizure events and note changes in network coherence over events without relating metrics back to clinically annotated EZ (33; 48). Other studies apply network methods, computing inter-electrode coherence, and relate these measures back to clinically annotated EZ or resection regions, but on data collected from a relatively small set of patients (31; 32; 47; 51). Studies that incorporate computational modeling to explain mechanisms of seizures and the EZ include (31; 51).

Here, we show a novel network-based algorithm that takes advantage of a certain type of signal evolution (ranked eigenvector centrality) and utilizes preictal, ictal and postictal data for tissue suspected to be within the EZ. Our study combines data from 4 centers and analyzes a total of 113 seizures from 42 patients. We compute network-based statistics and relate the eigenvector centrality (EVC) patterns back to clinically annotated EZ in patients with both successful and failed outcomes. We recently demonstrated that intracranial EEG (iEEG) is rich in network information beyond the typical signatures clinicians use to identify the EZ (12; 30; 46; 63). In particular, we modeled the epileptic brain as a dynamic networked system where EEG signals are correlated both temporally and spatially. We constructed a set of network-based statistics whose temporal evolution distinguishes the epileptic nodes from the non-epileptic nodes within specific epileptic networks, thus defining an electrophysiological signature of the EZ (30; 63). The electrophysiological signature of the EZ has a characteristic arch shape when visualized in a two-dimensional principal component (2D PC) space described below. The arch shape is significant because it indicates that the electrodes have lower centrality before a seizure, become highly central during a seizure, and then become less central after seizure offset. This suggests that the EZ is a brain region that becomes highly centralized when seizures occur, recruiting many other brain regions to participate in epileptic activity. We used these time series network-based statistics and the identified EZ arch signature to develop an algorithm that takes as inputs iEEG data and the patient’s brain image after electrode implantation and outputs the likelihood of an electrode being in the EZ.

We hypothesized that a network based-algorithm will show higher degrees of agreement with the clinically labeled EZ for successful surgical outcomes and lower degrees of agreement with the labeled EZ for failed surgical outcomes. Our hypothesis is based on our expectations that a network based-algorithm will perform favorably because epilepsy is a network disease of the brain and simply looking at biomarkers of individual electrodes ignores this fact. To test our hypothesis, we evaluated our algorithm in a blind, retrospective study on 42 patients that had undergone invasive monitoring and in most cases were followed by surgery. EEG data on 1-3 seizures was analyzed by our algorithm without knowledge of the seizure outcomes. Clinically identified EZ nodes were then compared to the most central nodes as defined by our algorithm. We found that the algorithm agreed more with clinical annotations for patients with successful surgical outcomes and less for patients with failed surgical outcomes. Since, HFO is considered a gold-standard for localization of high frequency power, we wanted to compare our results with such a method. We also applied qHFO algorithm presented in (18) to all patients whose EEG recordings met the requirements of the qHFO algorithm. We found that there were many patient datasets that could not be easily applied to the qHFO algorithm due to limitations on data available and sampling rates of equipment. However, on the datasets that could be compared with our network algorithm, there was a higher degree of agreement (DOA) with clinicians using a network algorithm versus only the qHFO algorithm.

Localization of the EZ is currently a time-consuming process since clinicians and technicians visually inspect fairly large data sets. In today’s data science era, it is important to develop and test computational tools to assist in localization of the EZ. An assistive computational tool would not only likely reduce extra-operative monitoring time in the EMU, thereby cutting medical costs and decreasing complications associated with invasive monitoring, but could also improve seizure freedom rates, especially in the more difficult to localize patients (i.e. non-lesional MRI patients). In addition, the underlying network-based algorithm that performs EZ detection favorably will further our understanding of the organization and dynamics of brain networks in epilepsy disease. Our results suggest that epilepsy changes how the different nodes in the brain are connected, and that diseased nodes are more likely to be highly central in the neuronal network and have a high centrality signature.

## METHODS - DATA COLLECTION

Patients included in this study were surgically treated for medically intractable seizures at four different centers: Johns Hopkins Hospital (JHH), National Institute of Health (NIH), the University of Maryland Medical Center (UMMC) and the Cleveland Clinic (CC). All patients included in this study underwent invasive pre-surgical monitoring with either subdural grid-and-strip arrays or stereotactic EEG depth electrodes for seizure localization or mapping of eloquent areas. Decisions regarding the need for invasive monitoring and the placement of electrode arrays were made independently of this work and solely based on clinical necessity. The research protocol was reviewed by the Johns Hopkins Institutional Review Board (IRB), the National Institute of Neurological Disorders and Stroke IRB, the University of Maryland Medical Center IRB, and the Cleveland Clinic IRB. The acquisition of data for research purposes was done with no impact on the clinical objectives of the patient stay. Digitized data were stored in a IRB-approved database compliant with Health Insurance Portability and Accountability Act (HIPAA) regulations (e.g. server hosted behind a firewall with sftp and ssh access).

At all four centers, as part of routine clinical care, up to three board-certified epileptologists marked, by consensus, the unequivocal electrographic onset of each seizure and the period between seizure onset and termination. The seizure onset was indicated by a variety of stereotypical electrographic features, which include, but were not limited to, the onset of fast rhythmic activity, an isolated spike or spike-and-wave complex followed by rhythmic activity, or an electrodecremental response. Concurrently with the examination of the EEG recordings, changes in the patients behavior were sought from the video segment of video-EEG recordings. For each patient, we combined surgical notes about the electrodes corresponding to resected regions and postoperative follow-up information about how the resection affected the patient’s seizures. The surgery was deemed a success and the resected area determined to include the EZ if, at least six months after surgery, a patient reported no seizures or could manage their epilepsy with medications. Failure was defined as the inability to localize the EZ at all, or if the patient continued to have seizures that were not manageable with medications after the resection.

iEEG recordings were acquired through subdural grid arrays, subdural strip electrodes, or depth-electrode arrays in various combinations as determined by clinical assessment for patients with temporal, occipital, or frontal lobe seizures. Subdural grids have 20-64 contacts per array and were used in combination with subdural strips with 4-8 contacts or depth arrays, thus having 80-116 recording electrodes per patient over all. Intracranial contact locations were documented by post-operative CT co-registered with a pre-operative MRI. Signals were acquired using continuous multi-channel iEEG recordings collected over 5 days on average (min.: 2 days; max: 10 days). Clinical monitoring lasted 5-10 days per patient and included 2-7 clinical seizures. Then clinicians clipped what they deemed clean sets of data and passed it through a secure transfer for the data analysis.

There were a total of 42 subjects analyzed retrospectively in this study: 7 from NIH, 20 from JHH, 7 from UMMC, and 8 from the Cleveland Clinic. There were 26 total successful surgeries and 16 total failed surgeries. The total number of electrodes per patient was 111.86 ± 23.89. The total number of electrodes used in analysis per patient (after removal of noisy/faulty channels, references, EKG, etc.) was 70.82 ± 24.84. The size of the clinically annotated EZ (# electrodes) was 8.05 ± 4.34. The onset age was 17.21 ± 13.48 years old, while all patients now are 34.68 ± 12.30 years old. The subject groups for each center are shown in Fig 2.

**Figure 2.**
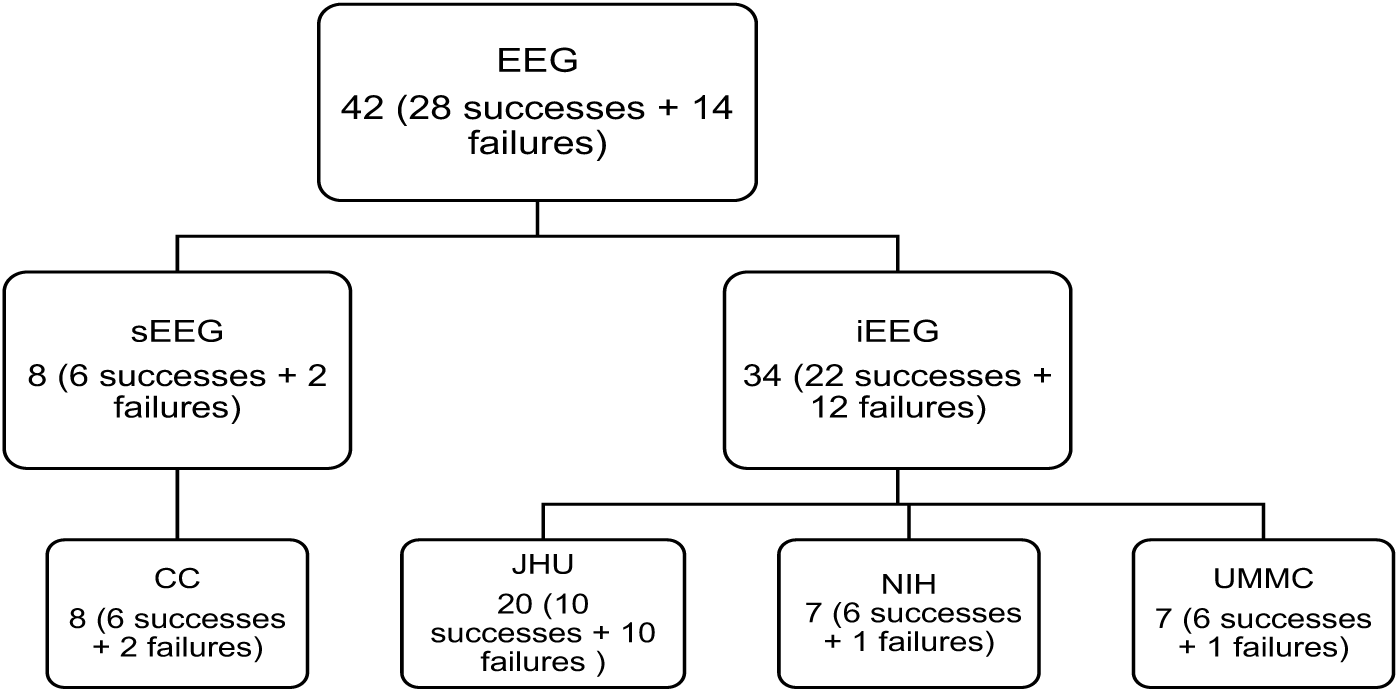
Patient cohort population for different recording systems, and across different hospital centers. Shows the distribution of successful and failed outcomes for each center.

### NIH Intracranial EEG Monitoring Technique - ECoG

Seven patients included in this study were surgically treated for drug-resistant seizures at the NIH NINDS and underwent invasive presurgical monitoring with subdural grids for seizure localization or mapping of eloquent areas. Recordings were acquired with a Nihon Kohden clinical EEG system. iEEG signals were sampled at a 1 kHz sampling rate and, filtered using a 300 Hz anti-aliasing filter. Signals were referenced to a common contact placed subcutaneously on the scalp, on the mastoid process, or on the subdural grid. Each data file stores continuous iEEG data from all channels and is automatically generated by the acquisition system.

### Johns Hopkins Hospital Intracranial EEG Monitoring Technique - ECoG

Twenty patients included in this study were surgically treated for drug-resistant seizures at the Johns Hopkins Hospital and underwent invasive presurgical monitoring with subdural grid and strip arrays for seizure localization or mapping of eloquent areas. Recordings were acquired with a Nihon Kohden clinical EEG system with a 1 kHz sampling rate and a 300 Hz anti-aliasing filter, and were converted to EDF format for storage and further processing. Each EDF file stores approximately 42 minutes of continuous ECoG data from all channels and is automatically generated by the acquisition system. Consecutive EDF files cover consecutive, non-overlapping, time windows with less than 5s-lag in between. Digitized data were stored in a IRB-approved database compliant with HIPAA regulations.

### UMMC Intracranial EEG Monitoring Technique - ECoG

Seven patients included in this study were surgically treated for drug-resistant seizures at the University Maryland School of Medicine and underwent invasive presurgical monitoring with subdural grid and strip arrays for seizure localization or mapping of eloquent areas. At the University of Maryland Medical Center (UMMC), recordings were acquired with a Natus/XLTEK system (Natus Medical Incorporated, Inc., Pleasanton, CA) with 250-1000 Hz sampling rate and 50-300 Hz anti-aliasing filter, and were converted to EDF format for storage and further processing. Each EDF file stores approximately 42 minutes of continuous ECoG data from all channels and is automatically generated by the acquisition system. Consecutive EDF files cover consecutive, non-overlapping, time windows with less than 5s-lag in between. Digitized data were stored in a IRB-approved database compliant with HIPAA regulations.

### Cleveland Clinic Stereotactic EEG Monitoring Technique - SEEG

Eight patients that underwent SEEG invasive monitoring from the Cleveland Clinic epilepsy center were included in this study. The choice of electrode location was based on a pre-implantation patient management conference and was made independently of the present study. Criteria for patients undergoing SEEG implantation were reviewed by clinicians to determine patient eligibility for enrollment in the current study. If the patient met study criteria, research staff not involved in the surgery implantation or post-surgical care contacted the patient for potential participation in the study.

For each subject, approximately 8-13 stereotactically placed depth electrodes were implanted. The electrode contacts were 0.8 mm in diameter, 2 mm in length, and spaced 1.5 mm apart. Depth electrodes were inserted in either orthogonal or oblique orientations using a robotic surgical implantation platform (ROSA, Medtech Surgical Inc., USA) allowing intracranial recording from lateral, intermediate and/or deep cortical and subcortical structures in a three-dimensional arrangement (21). The day prior to surgery, volumetric pre-operative MRIs (T1, contrasted with Multihance 0.1 mmol/kg) were obtained and used to pre-operatively plan electrode trajectories. All trajectories were evaluated for safety; any trajectory that appeared to compromise vascular structures was adjusted appropriately without affecting the sampling from areas of interest.

SEEG electrophysiological data was acquired using a conventional clinical electrophysiology acquisition system (Nihon Kohden 1200, Nihon Kohden America, USA) at a sampling rate of 1 kHz and 300 Hz anti-aliasing filter. Behavioral event data were simultaneously acquired during behavioral experiments along with the SEEG electrophysiology and stored for subsequent analysis. All signals were referenced to a contact affixed to the skull. Archived electrophysiological data was not filtered prior to offline analysis.

Each patient had electrode contacts characterized according to anatomical location. The anatomical locations of all contacts were identified through inspection of post-operative imaging, requiring agreement by two clinical experts. An example of post-operative imaging contributing toward determining contact location is shown in 1. Coronal and sagittal views were available for every contact.

## METHODS - COMPUTATIONAL STEPS

In this study, our raw dataset consisted of EEG recordings of seizures with 60 seconds of data before and after each seizure. Data was collected from 42 patients with at least two seizures per patient. We applied network analysis techniques and considered each electrode in the iEEG array to be a node in a network. The overall process of our algorithm is highlighted in Fig. 3. We computed the cross-power spectrum matrix for each time window, then the corresponding EVC and then we trained a Gaussian weighting function that assigned a likelihood to each electrode for being within the EZ. After computing the heat map for the EZ predicted set of electrodes, we compared them to the clinical electrodes for both successful and failed surgical outcomes. We show results for each center separately, and also all patients grouped together. Note that we trained the Gaussian weighting function only using one center’s patients, so that we could test our results across center. Clinical procedures can vary more from center to center versus the variability within center, so it is a conservative approach to train using one center and then test on all other centers to see if our analysis holds across different clinical procedures.

**Figure 3.**
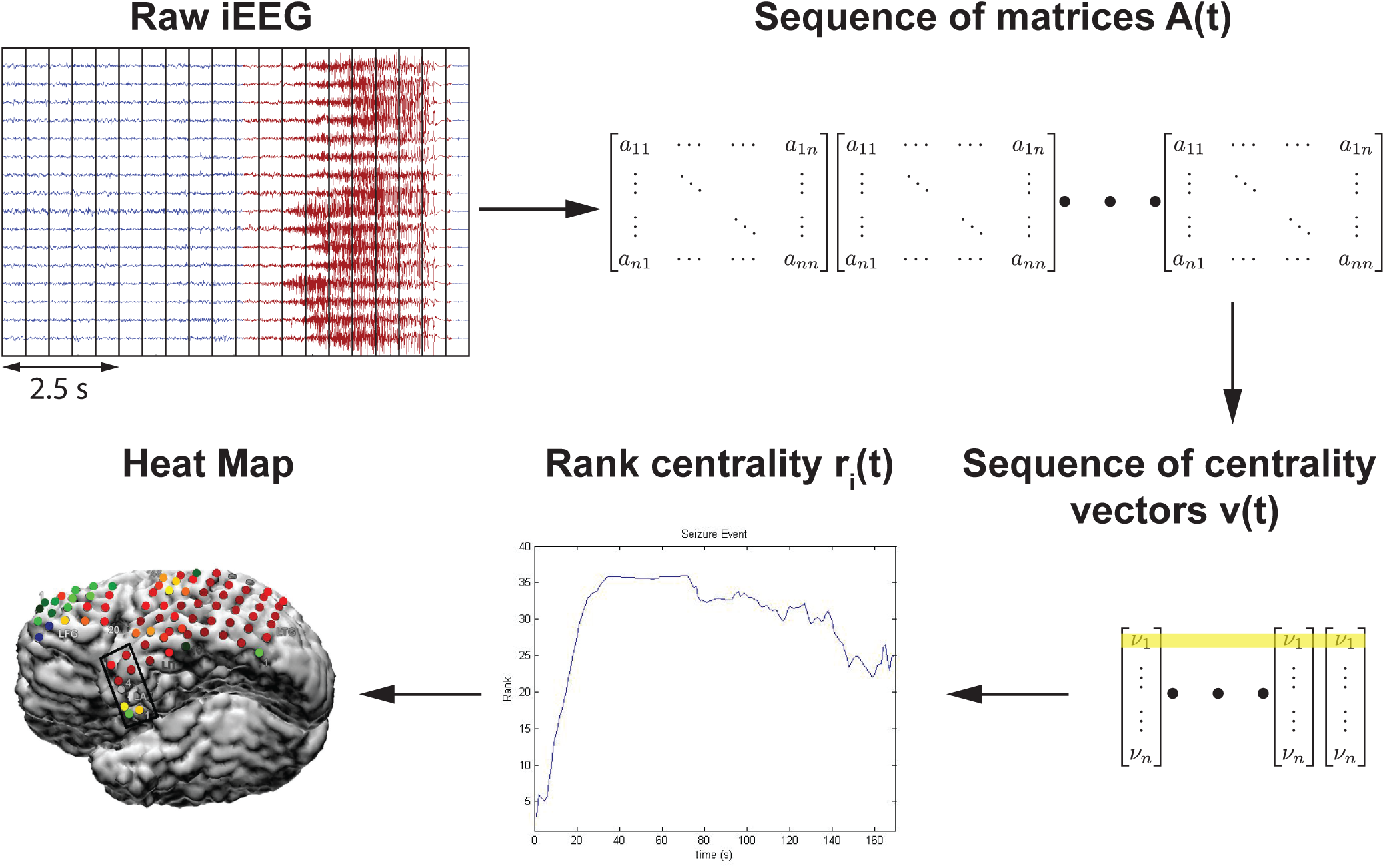
Computational steps for seizure onset localization: the algorithm processes raw ECoG to compute the sequence of adjacency matrix A(t). From this sequence, A(t), it computes the sequence of leading eigenvectors, v(t), as a network centrality measure, the EVC. Algorithm then converts EVC into the sequence of rank centrality r(t). From this sequence, r(t), algorithm computes a heatmap that generates predictions of the EZ. Yellow shading indicates the EVC of 1^st^ electrode evolving in time whose rank centrality, *r_1_(t)*, is illustrated in the plot.

All Matlab (R2016b) and Python (v 2.7) code is publicly available online at: https://github.com/ncsl/eztrack.

### Preprocessing of Data

All data underwent digital filtering with a butterworth notch filter of order 4, implemented in MATLAB with the *filtfilt* function (frequency ranges of 59.5 to 60.5). In general, EEG data is known to be noisy and referencing schemes can play a significant role in downstream data analysis. We decided to apply a common average referencing scheme to the data before analysis (37). Here, we take an average signal from all recording electrodes and subtract it from the electrodes. This has been shown to produce more stable results and rejects correlated noise across many electrodes (18). We made sure to exclude any electrodes from subsequent analysis if they were informed to have artifacts in their recording by clinicians.

### Compute and Rank Nodal Centrality Over Time

Network centrality for each node was computed every second using a 2.5 second sliding window sliding every second 60 seconds before seizure, during seizure, and 60 seconds after seizure for at least 2 seizure events. For each window, the brain network was first represented by a connectivity matrix (15), by computing all pairwise cross-power spectra between the signals in the gamma frequency band (30-90 Hz), i.e.,

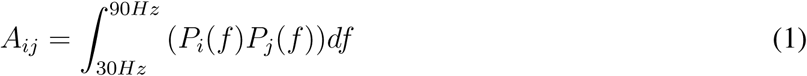

where *P_i_*,*P_j_* are the magnitudes of the Fourier transform of the time series in the window recorded from electrodes *i*, *j*, and A_*ij*_ is the element of connectivity matrix and is the adjacency between nodes *i* and *j*. We chose the gamma band because the gamma frequency band has often exhibited the most modulation in power between non-seizure and seizure periods. It has been thought to be correlated to neuronal spiking and fMRI activity and thus carries information in such invasive recordings (22; 61; 62).

The importance of each electrode to the network connectivity was measured by the strength and number of connections it makes with other electrodes, referred to as centrality. We used the eigenvector centrality (EVC) to measure the connectivity of each electrode, as EVC showed interesting repeatable patterns over seizure events in our prior study (12). The EVC of an electrode is defined as the sum of the EVCs of all other electrodes weighted by their connectivity, which measures the relative influence of a node within the network. The EVC of all electrodes is computed implicitly as:

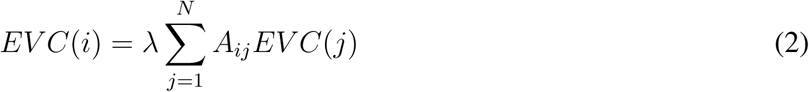

λ is the leading eigenvalue of the connectivity matrix A and the EVC is then the leading eigenvector of A. In simple terms, the EVC of a node in the network (electrode) is proportional to the sum of EVCs of its neighbors (nodes it is connected to). That is, a node is important if it is (i) connected to a few nodes that are themselves very important or if it is (ii) connected to a very large number of not-so-important nodes. The leading eigenvectors of connectivity matrices were calculated numerically at each second during the recordings from the connectivity matrices. Finally, the EVC vector for each second was converted to a ranked vector containing values 1 to *N*, where a 1 was placed in the component of EVC that had the smallest centrality and an N was placed in the component of EVC that had the largest centrality.

### Normalize Rank Evolution Signals

Next, we normalized the rank evolution signals (the EVC) for each electrode in the *X* (time) and *Y* (rank centrality, i.e. number of electrodes) directions. This was done so that we can compare signals from different patients that have varying numbers of electrodes and varying seizure durations across individuals and within individuals. To normalize along the *X*-axis, we either stretched (interpolated) or shrunk (simply downsampled at a lower sampling rate) each ranked EVC signal during a seizure epoch such that all signals were 500 data points in length. Most ranked EVC signals were under 500 seconds in length, so the majority of the rank centrality signals were stretched using linear interpolation (using the interp1 function in Matlab) preserving the shape of the signal during a seizure event. To normalize along the *Y*-axis, we scaled the rank centrality between 0 and 1 by dividing by the number of electrodes. Further, in order to compare the ranked EVC in a quantifiable manner, we normalized all the *X*, *Y* normalized signals such that the centrality signal integrated to 1. We divided the normalized rank centrality by area under the curve. This normalization converted each signal into a probability density function,

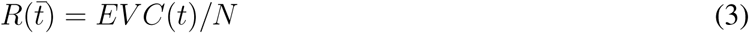

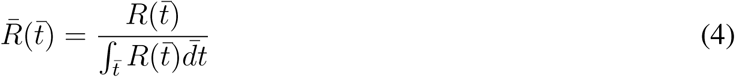

where 
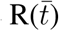
 is the normalized rank signal in time after dividing by the number of electrodes and 
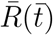
 is the normalized rank signal at normalized time 
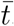
.

### Compute Feature Vector from Normalized Rank Signals

For each normalized signal, we extracted the deciles in time, the locations at which the signal integrates equally to 10% of the total area under the curve, i.e. points in normalized time where the signal integrates to 0.1, 0.2, 0.3, and so on until the end of the signal is reached. This gives a 10 dimensional vector for each signal that serves as a *feature vector*.

### Electrode Weight Assignment Based on Feature Vectors

Once we calculated feature vectors for each signal, we projected the features into a 2D principle component (PC) space. This was done by assuming that each feature vector is an observation, hence the analysis was performed in *space x time*. We performed PC analysis and plotted the features across all electrodes and patients projected onto the first and second PCs. Each electrode (data point in Fig. 4A) was labeled according to whether or not the electrode was in the clinical annotated EZ region and whether the surgical resection was a success or a failure. We then created a weighting function over the 2D PC space, which would assign a weight to an electrode based on their location in PC space.

To generate this weighting function, we discretized it into equally sized square partitions (100 × 100 along 1^*st*^ and 2^*nd*^ principal components). The mean normalized rank signature across all data points was computed for each partition. The signatures for the four corner partitions are shown in Fig. 4A. The shapes of the mean normalized rank signatures across partitions change in a somewhat continuous manner. Moving vertically from the bottom of the PC space to the top, the rank signatures transition from a concave to a convex shape. Moving from left to right, the signature shifts horizontally: forward (to the right) if the partition is at the bottom of the PC space, and backwards (to the left) if at the top of the PC space.

Our hypothesis is that the arch signature displayed in the bottom left of Fig. 4A represents the signatures of the EZ because this is the region of the PC space that has the most isolated channels that come from patients with successful outcomes (green + points). In fact, the bottom portion of the PC grid shows the arch signature. Therefore, the weighting function is set to be highest in these regions and decay as a function of distance from these regions. We defined a weighting function to be the sum of 4 bivariate Gaussian-like functions (Eq. 5, Fig. 4B) as shown in (5). The 2D PC space is divided into 4 quadrants defined by an origin. See Fig. 4B (left) with origin (−100, –100).

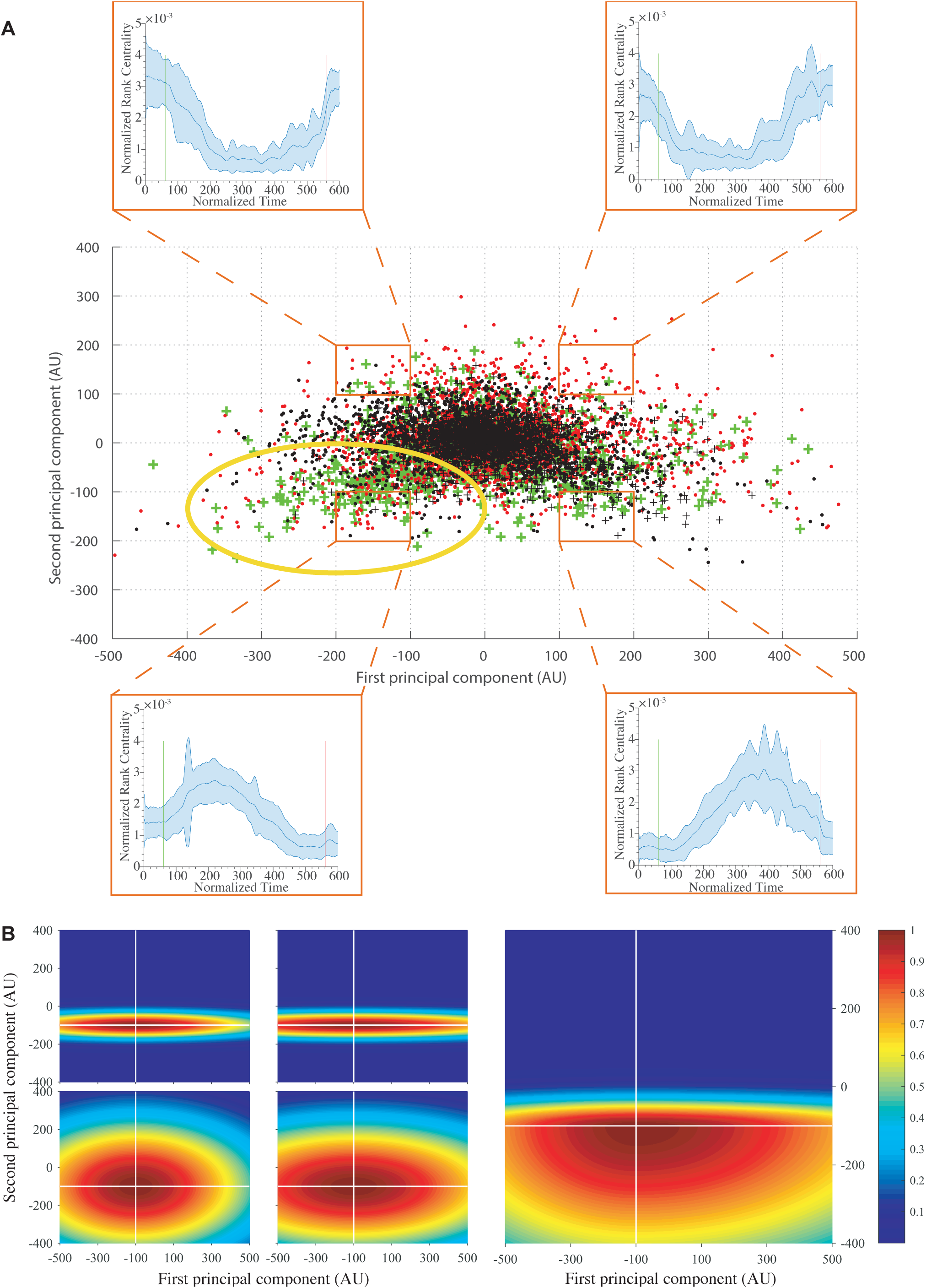

### Training Origin of Gaussian Weighting Function

In each quadrant, the bivariate Gaussian-like function were initialized with the shapes in Fig 4A. The covariance matrix in each quadrant was computed as the sample covariance from the data points in that quadrant. The origin of the four quadrants is the mean vector, which is trained. We followed a leave-one-out training procedure on the sample of 20 patients collected at JHU. We chose JHU because it had the greatest number of patients collected within center and would still account for less then 50% of the total patients. The mean of all four quadrants is optimized for maximizing the DOA. In Fig 4B, this is shown as (−100; 100), which was found at the end. Once the optimized mean is found, then all four quadrant’s Gaussian functions, *w_i_(x,y)*, are linearly combined with a heaviside step function to get the final Gaussian weighting function, *w(x,y)*. This final Gaussian weighting function, *w(x,y)* is used to assign weights to all subsequent EVC of each electrode for every patient. This in turn produces the likelihood of every electrode being within the EZ set.

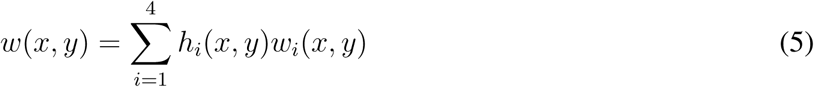

where

- 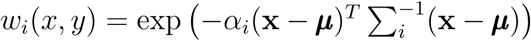
- *α*_*i*_ - exponential decay factor for *i*^*th*^ quadrant
- 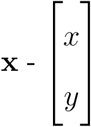
, and 
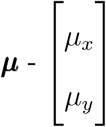
 define the position and mean vector respectively
- Σ_*i*_ - covariance matrix of *i*^*th*^ quadrant
- 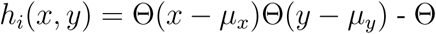
 is the heaviside step function
- *(x,y) ∈ i^th^* quadrant

### Computing Degree of Agreement and Statistical Analysis

For every seizure event for every patient in NIH, UMMC and CC, we generated a set of electrodes with their heatmap (defined by electrode weights; see Fig. 3), which can be interpreted as their likelihood for being in the EZ. For each seizure recording, we then computed the degree of agreement between the computed EZ likelihoods and clinical annotations of the EZ. The likelihood was computed using the Gaussian weighting function trained as described in the previous subsection. Then, a threshold α = 0.3, 0.6, 0.9 was applied to each heatmap and the set of electrodes whose likelihoods exceeded α were defined as the algorithm’s EZ (AEZ). The AEZ was then compared to clinically annotated EZ (CEZ) using the following degree of agreement (DOA) statistic:

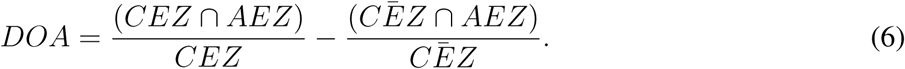

Note that 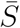
 is the complement of the set *S*, and that *D* ∈ [−1,1], where *DOA* = 1 implies perfect agreement and DOA < 0 is less agreement.

Across all patients, electrodes, and seizure events, we have a collection of DOA values. We then derive two distributions: (i) the distribution of DOA for all electrodes implanted in patients who had successful treatments, and (ii) the distribution of DOA for all electrodes implanted in patients who had failed treatments. We then test whether there is a significant difference in DOA distribution between these two patient groups using the Wilcoxon rank sum test to test for statistical differences. This non-parametric test was selected, as the data are not guaranteed to meet the normality conditions for a Student’s t-test (58). In addition, we also added an across-center analysis where we combine all the data and test whether the DOA distributions for successful versus failed outcomes are significantly different.

On top of this analysis, we also add a minmax scaling to normalize the of degree of agreements within each center, so that success and failure could be compared at the same scale.

### High Frequency Oscillator - qHFO Detector

We compared our algorithm with the qHFO algorithm presented in (18), which uses a sensitive HFO detector, then redacts HFOs that were produced by artifacts. Previous work has shown that sampling rates of 1000 Hz are capable of recording HFOs, but only capture 60% of the events (18). Therefore, we only analyzed patients with sampling rates ≥ 1000 Hz and with available interictal data. This resulted in 3 patients from NIH and 2 patients from JHU, with a total of 13 separate recorded datasets. The datasets here analyzed had an average recording of 7.1 min, 83 total electrodes analyzed, and 10 electrodes within the clinically annotated EZ set. Using the qHFO algorithm on this data required a few minor adaptions.

We used a single common average reference applied to all analyzed intracranial electrodes (as described earlier), rather than separating the referencing between depth electrode channels and grid channels as was done in (18). The popDet artifact rejection method also could not be used, as it requires sampling rates of at least 2,000 Hz.

## RESULTS

Every patient (n=42) with at least two seizures was analyzed (total of 113 seizures) with 20 of the patients from JHU used to train the final Gaussian weighting function. The output of the process for each seizure recording is each electrode’s likelihood of being in the EZ. These likelihood scores are in turn used to produce a heatmap that can be overlaid on a brain MRI to show the relative predicted EZ region for a certain patient. Figure 5 shows a few examples of heatmaps for 3 patients who had successful outcomes and 3 patients with failed outcomes. For the 3 successful patients, the AEZ lies entirely within the resected regions, suggesting a high DOA between the AEZ and CEZ. For one of the failed patients, the resected region and the AEZ do not overlap, i.e. DOA is low. For the other failed patient, the AEZ is a very small set, suggesting that the EZ may not be appropriately covered by the electrode implantation.

**Figure 5.**
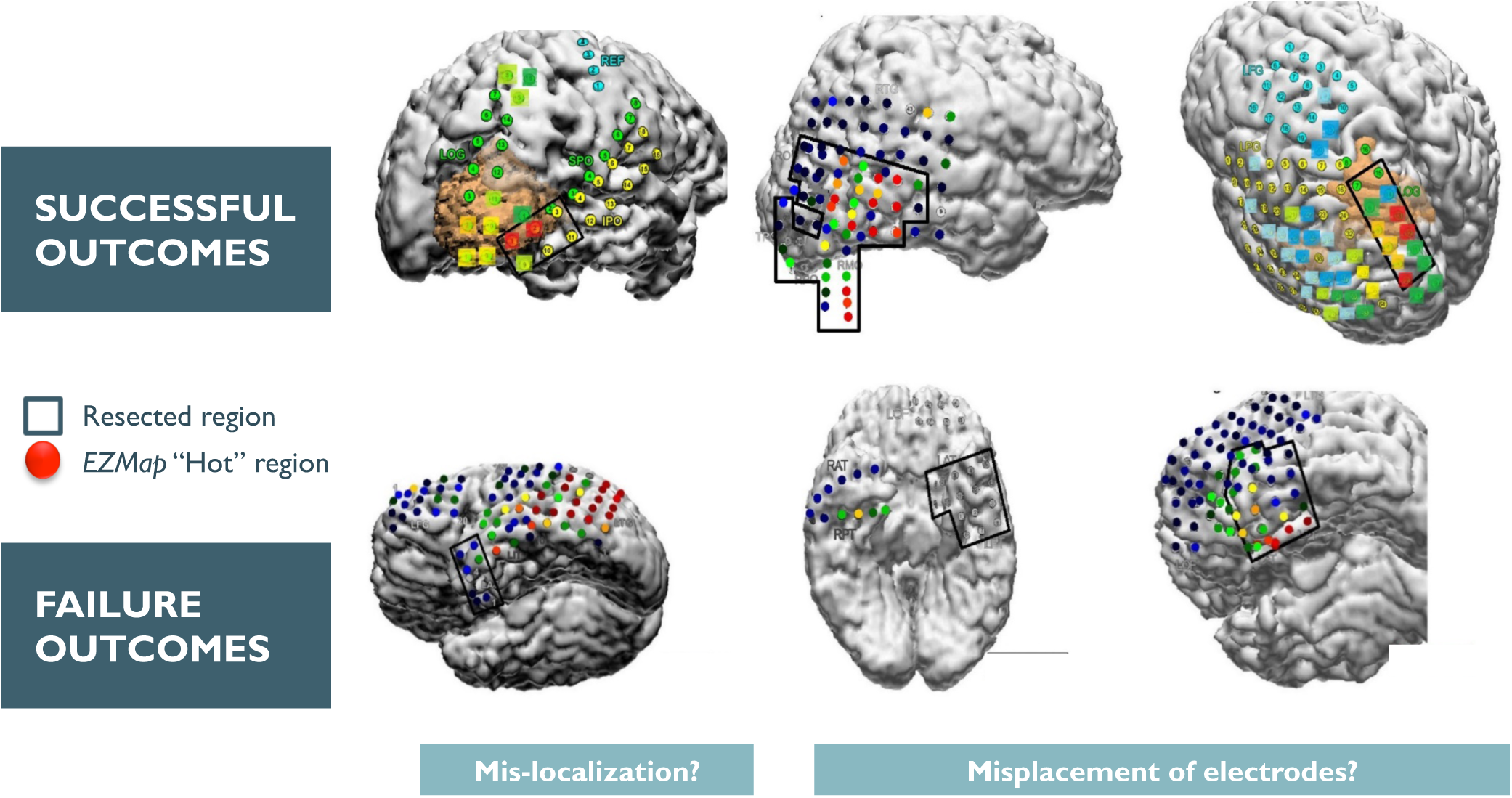
This shows an example overlay of the algorithm’s heatmap of likelihood on a brain scan for 6 patients (3 successful and 3 failed outcomes). The red region shows our predicted onset zone and the black outlines represent where the clinicians performed a resection. The orange, yellow, green and blue regions represent lower likelihoods for that specific electrode being within the EZ set as predicted by the algorithm.

In our comparative HFO analysis, we analyzed 13 segments of data from 5 patients. Of the 13 files, most patients have no HFOs, even at 1000 Hz sampling rate (see table 1). Only 3 data segments had HFO detections, but one of them did not have an anomalous grouping suggestive of the EZ (30% of the total recording time from all 13 data segments). In JH3, there were HFOs, but no channels had an anomalous rate high enough to be predicted within the EZ set. In NIH pt1aw2 and pt3alsp3, both only had a single channel predicted to be in the EZ. This prediction was in concordance with clinically annotated EZ in pt1 but not in pt3.

**Table 1.** HFO results for the 2 patients with interictal data from NIH. Only 2 datasets (2 patients) showed HFO rates not identically zero. Only 1 dataset had an HFO analysis with an electrode within the clinically annotated set.

| Patient | Duration (seconds) | Identification by HFO |
| --- | --- | --- |
| JH1 | 1800 | Rates identically zero |
| JH3 | 1800 | No anomalously high channels |
| pt1aslp1 | 405 | Rates identically zero |
| pt1aslp2 | 498 | Rates identically zero |
| pt1aw1 | 425 | Rates identically zero |
| pt1aw2 | 414 | Prediction has been made 'AD1' |
| pt2aslp1 | 376 | Rates identically zero |
| pt2aslp2 | 419 | Rates identically zero |
| pt2aw1 | 397 | Rates identically zero |
| pt2aw2 | 664 | Rates identically zero |
| pt3aslp1 | 362 | Rates identically zero |
| pt3aslp2 | 379 | Prediction has been made 'SFP6' |
| pt3aw1 | 363 | Rates identically zero |

The lower sampling rate and short time segments are not ideal for automated HFO analysis, as is apparent from these results. In our network analysis, we had a high DOA with pt1 (0.62), while a relatively lower DOA for pt3 (−0.16). It seemed that for pt3, HFO analysis completely disagreed with clinical annotations, while the network analysis found more electrodes then the clinically annotated EZ, which led to lower DOA. For pt1, the network analysis also highlighted the same electrode as being in the EZ set. This shows how HFO and network analysis can complement each other in analyzing different sections of the data. Based on our limited comparisons due to inherent data limitations, our analysis is more capable of identifying the full clinically annotated EZ then HFOs in this specific dataset.

In Fig. 6, we show the degree of agreement (DOA) for datasets collected from the test datasets (the three clinical centers: UMMC, NIH, CC) for 3 different threshold values, α, that is placed on the likelihood distribution (electrodes with likelihood greater then threshold are placed in EZ set). The resulting DOA after training the Gaussian weighting function for JHU are shown in supplementary information. It also shows the same trend as seen in Fig. 6. As illustrated in Fig. 6, the general trend is that the DOA distributions for successes and failures separate more as increases, and α = 0.9 appears to be an operative threshold that shows a positive DOA for successes and a negative DOA for failed outcomes. For α = 0.9, the statistics for DOA (mean and stdev) are given in Table 2 for each center and across all centers together. By applying a Wilcoxon rank-sum test, we also see a significant difference at significance level 0.05 for all centers at threshold level of 0.9. At each center, there is a trend of the DOA that is a function of clinical outcome of the patient. This is consistently shown across recording platform (ECoG for UMMC, NIH and SEEG for CC) and patient population. In all cases, as the threshold increases from 0.3 to 0.9, the difference of DOA between success and failed cases increases. If there is low DOA with the algorithms EZ and the clinically annotated EZ and the patient is a failed outcome, then this may be a case of mislocalization. If, on the other hand, there is no visible EZ from the algorithm (all weights are low), then the EZ may not be in the vicinity of the electrode, suggesting a possible misplacement of electrodes.

**Table 2.** Degree of Agreement Results for α = 0.9 with average ± standard deviation from each clinical center and also the resulting p value from the Wilcoxon ranked sum test. All centers show a significant difference between success and failure cases. Note JHH is used in the training of the Gaussian weighting function.

| Center | DOA Statistics for Success | DOA Statistics for Failure | P Value |
| --- | --- | --- | --- |
| UMMC | $0.09 \pm 0.15$ | $-0.09 \pm 0.08$ | 0.027 |
| NIH | $0.21 \pm 0.25$ | $-0.32 \pm 0.11$ | 0.020 |
| CC | $0.01 \pm 0.38$ | $-0.38 \pm 0.01$ | 0.024 |
| *JHH | $0.21 \pm 0.23$ | $0.08 \pm 0.25$ | 0.016 |
| All | $0.14 \pm 0.27$ | $0.00 \pm 0.27$ | 0.002 |

**Table 3.** Degree of Agreement Results for α = 0:9 with average ±standard deviation from each clinical center after minmax scaling and also the resulting p value from the Wilcoxon ranked sum test. All centers show a significant difference between success and failure cases.

| Center | DOA Statistics for Success | DOA Statistics for Failure | P Value |
| --- | --- | --- | --- |
| UMMC | $0.35 \pm 0.23$ | $0.05 \pm 0.11$ | 0.0057 |
| NIH | $0.54 \pm 0.23$ | $0.08 \pm 0.07$ | 0.0061 |
| CC | $0.36 \pm 0.34$ | $0.02 \pm 0.01$ | 0.0016 |
| *JHH | $0.50 \pm 0.23$ | $0.36 \pm 0.24$ | 0.0158 |
| All | $0.45 \pm 0.27$ | $0.29 \pm 0.25$ | 0.0005 |

**Table 4.** Table of patient data for each center describing patient characteristics and electrode statistics. Some data was not available clinically and is represented by N/A.

| Years | Sampling Freq (Hz) | Gender | Hand Dominant | Center | Total # Electrodes | # Elec Used (Analysis) | Size of Clinical EZ (# elec) | Strip Electrodes Present? (y/n) | Outcome (S/F) |
| --- | --- | --- | --- | --- | --- | --- | --- | --- | --- |
|  | 1000 | F | R | CC | 120 | 94 | 4 | n | F |
|  | 1000 | F | R | CC | 120 | 79 | 4 | n | S |
|  | 1000 | M | R | CC | 150 | 111 | 19 | n | S |
|  | 1000 | F | R | CC | 90 | 78 | 4 | n | S |
|  | 1000 | M | R | CC | 110 | 93 | 6 | n | F |
|  | 1000 | M | R | CC | 70 | 46 | 4 | n | S |
|  | 1000 | F | R | CC | 130 | 99 | 6 | n | S |
|  | 1000 | M | R | CC | 90 | 67 | 2 | n | S |
|  | 500 | M | Right | UMMC | 128 | 88 | 3 | n | F |
|  | 500 | M | Right | UMMC | 128 | 49 | 10 | y | S |
|  | 250 | M | N/A | UMMC | 128 | 45 | 10 | y | S |
|  | 250 | M | Right | UMMC | 128 | 46 | 6 | y | S |
|  | 1000 | M | N/A | UMMC | 128 | 47 | 11 | y | S |
|  | 250 | M | N/A | UMMC | 128 | 52 | 9 | y | S |
|  | 1000 | M | N/A | UMMC | 128 | 30 | 17 | y | F |
|  | 1000 | F | R | NIH | 98 | 86 | 10 | n | S |
|  | 1000 | F | R | NIH | 86 | 64 | 8 | n | S |
|  | 1000 | M | R | NIH | 135 | 99 | 12 | n | S |
|  | 1000 | M | L | NIH | 80 | 59 | 11 | n | S |
|  | 1000 | F | R | NIH | 89 | 53 | 9 | n | F |
|  | 1000 | F | R | NIH | 65 | 54 | 6 | n | F |
|  | 1000 | M | R | NIH | 132 | 119 | 6 | n | S |
|  | 1000 | N/A | N/A | JHU | N/A | N/A | N/A | N/A | F |
|  | 1000 | N/A | N/A | JHU | N/A | N/A | N/A | N/A | S |
|  | 1000 | N/A | N/A | JHU | N/A | N/A | N/A | N/A | F |
|  | 1000 | N/A | N/A | JHU | N/A | N/A | N/A | N/A | S |
|  | 1000 | N/A | N/A | JHU | N/A | N/A | N/A | N/A | F |
|  | 1000 | F | N/A | JHU | N/A | N/A | N/A | N/A | S |
|  | 1000 | F | N/A | JHU | N/A | N/A | N/A | N/A | F |

**Table 5.**
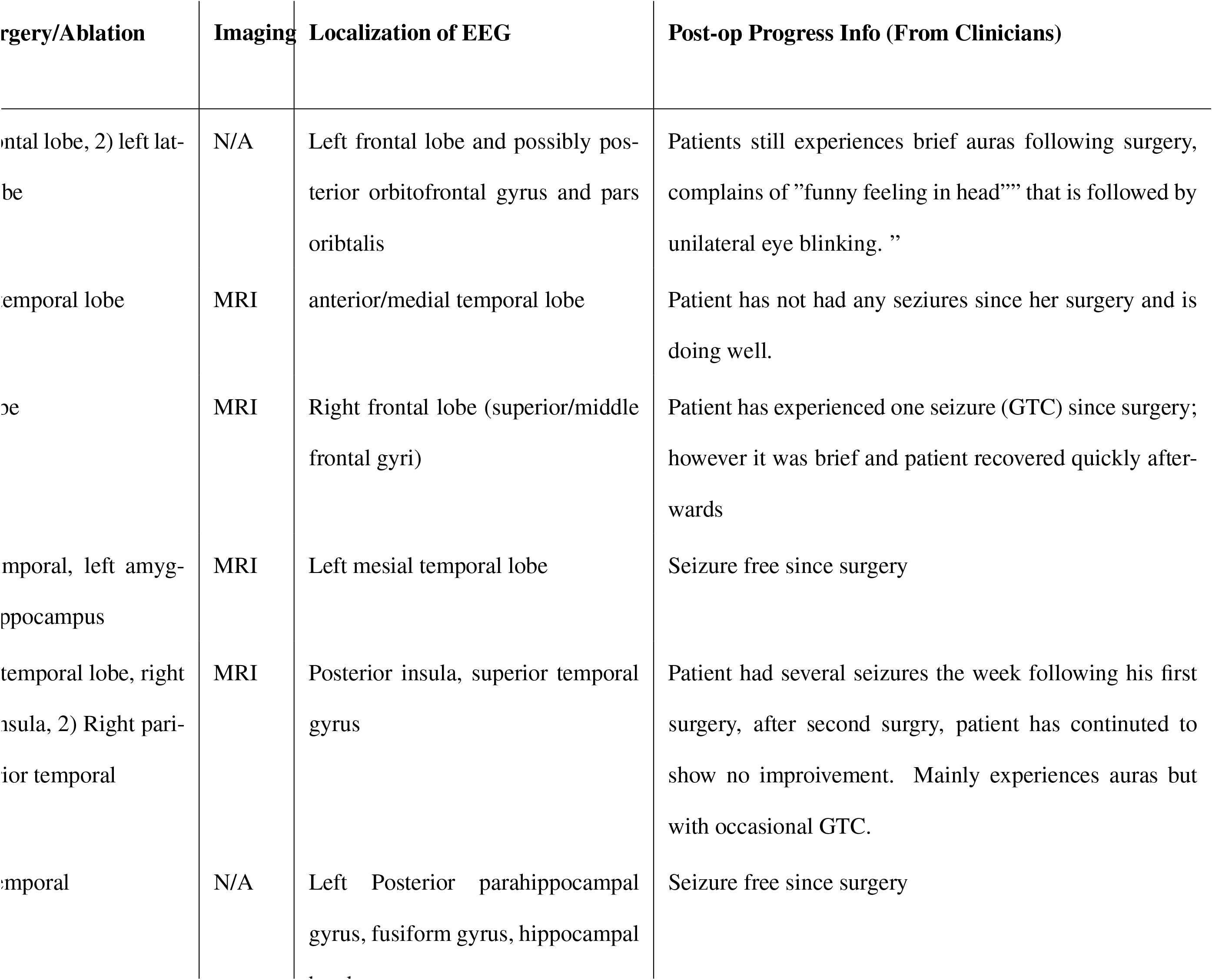
Table of clinical notes for each patient at Cleveland Clinic from their clinical procedures (imaging, resection). Some data was not available clinically and is represented by N/A.

**Table 6.**
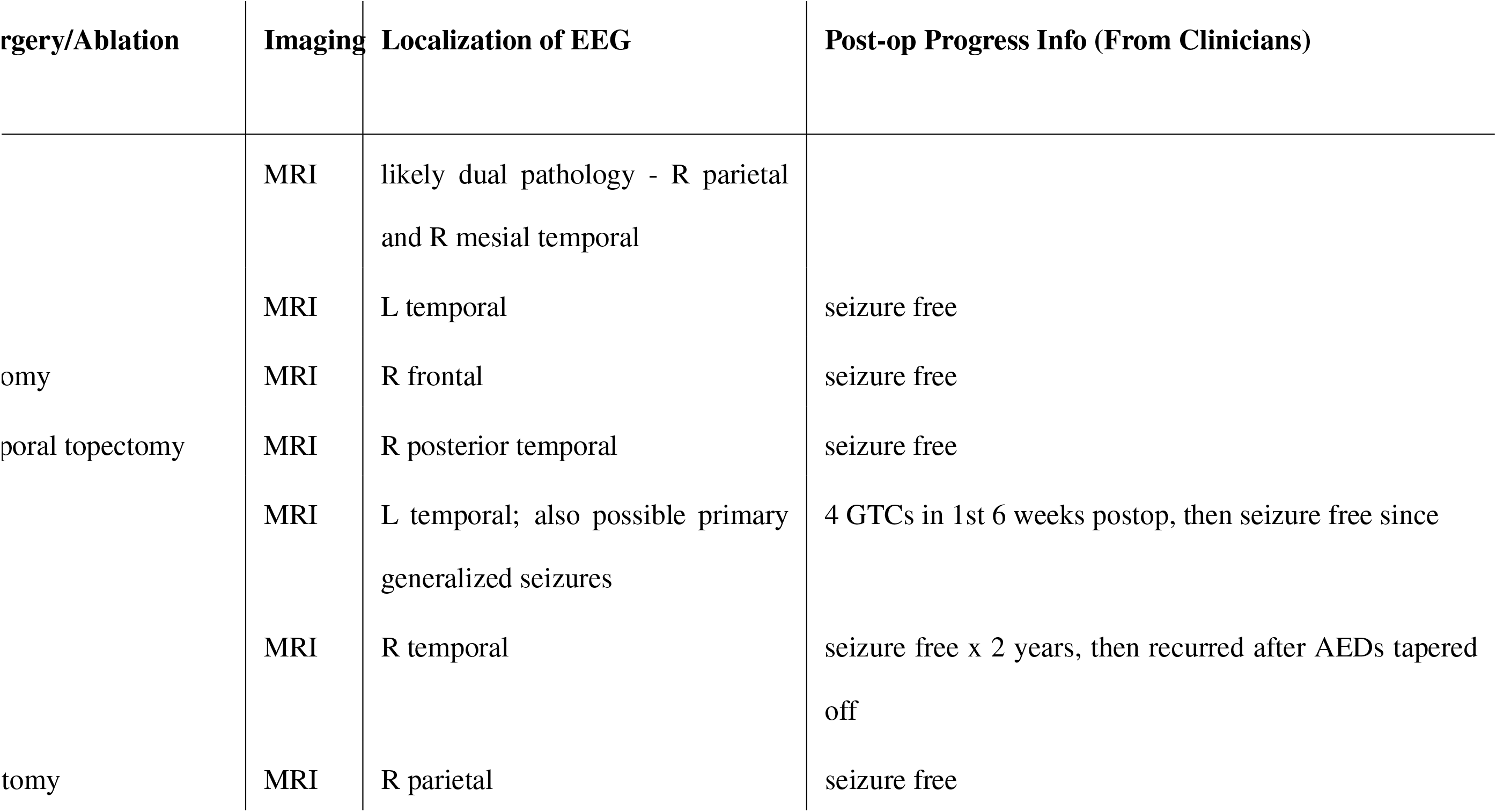
Table of clinical notes for each patient at NIH from their clinical procedures (imaging, resection). Some data was not available clinically and is represented by N/A.

**Table 7.**
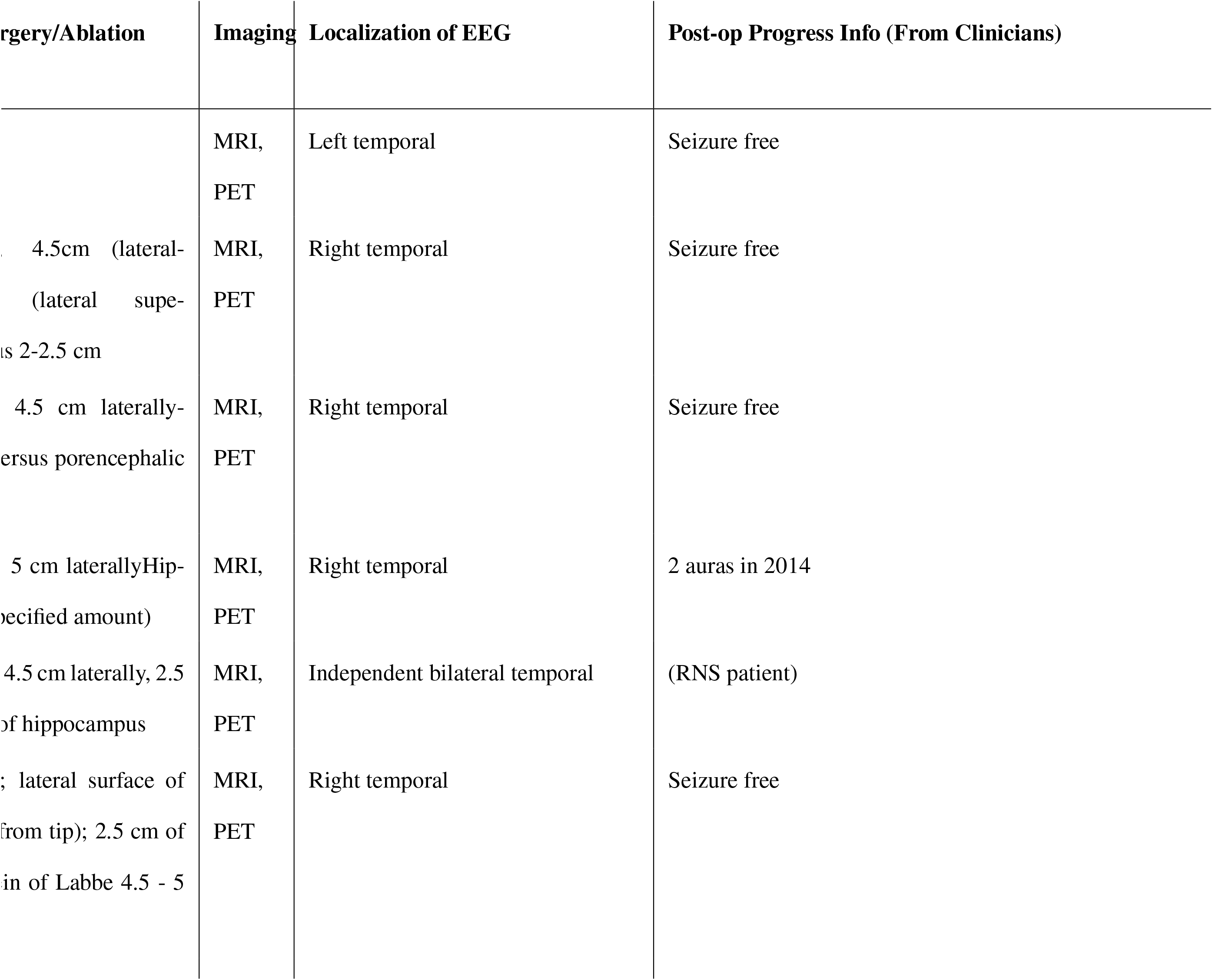
Table of clinical notes for each patient at UMMC from their clinical procedures (imaging, resection). Some data was not available clinically and is represented by N/A.

**Table 8.**
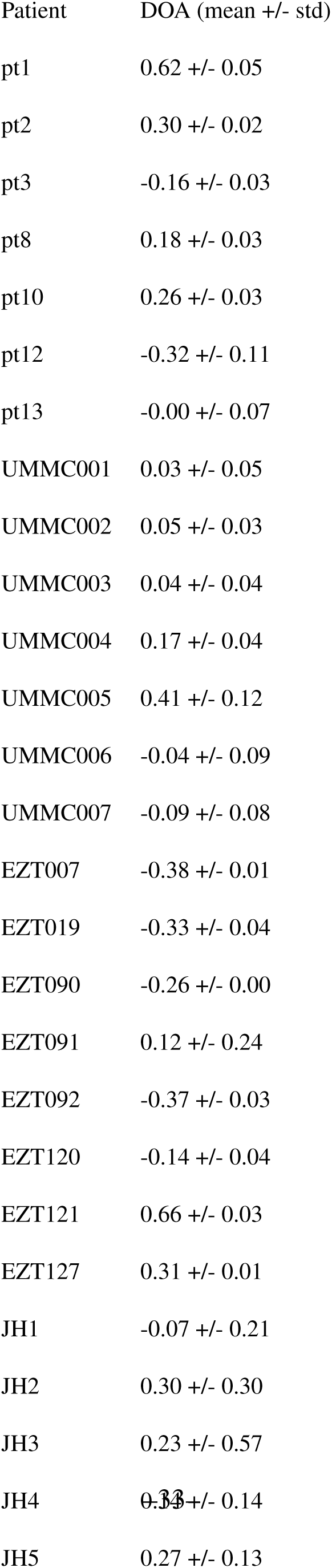
Table of DOA scores for each patient separately. Each patient has 2-3 seizure recordings available and a DOA score was computed for each recording instance. The JHU scores are also included after the end of the leave-one-out procedure.

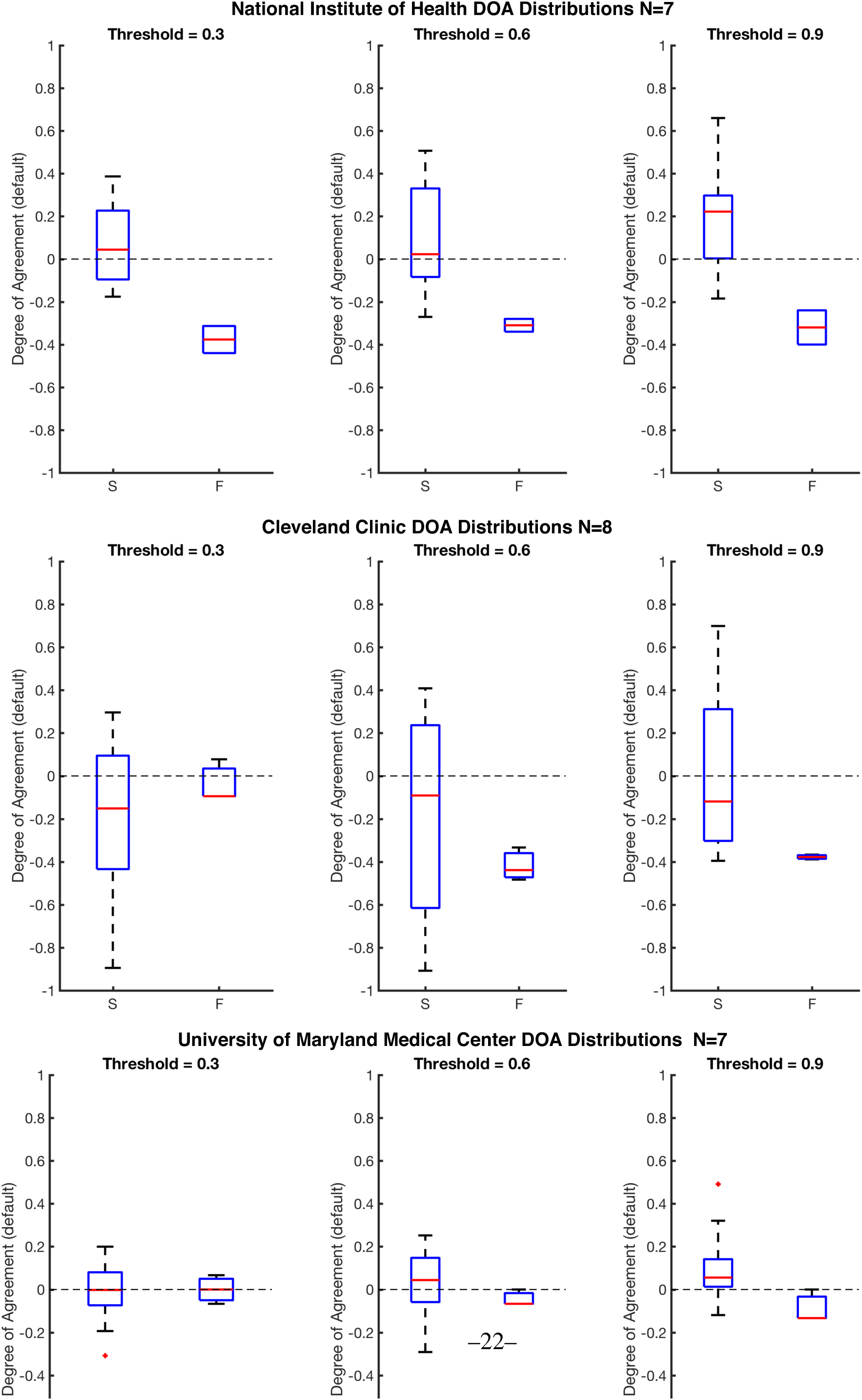

We also show in Figure 7 that there is no bias due to center (reulsts are shown in 3). All centers, when normalized show a significant difference between success and failures. The large variation is due to the varying number of electrodes implanted per patient and the varying size of the clinical EZ hypothesis. However, all centers show significant difference when compared with a Wilcoxon Ranksum test.

**Figure 7.**
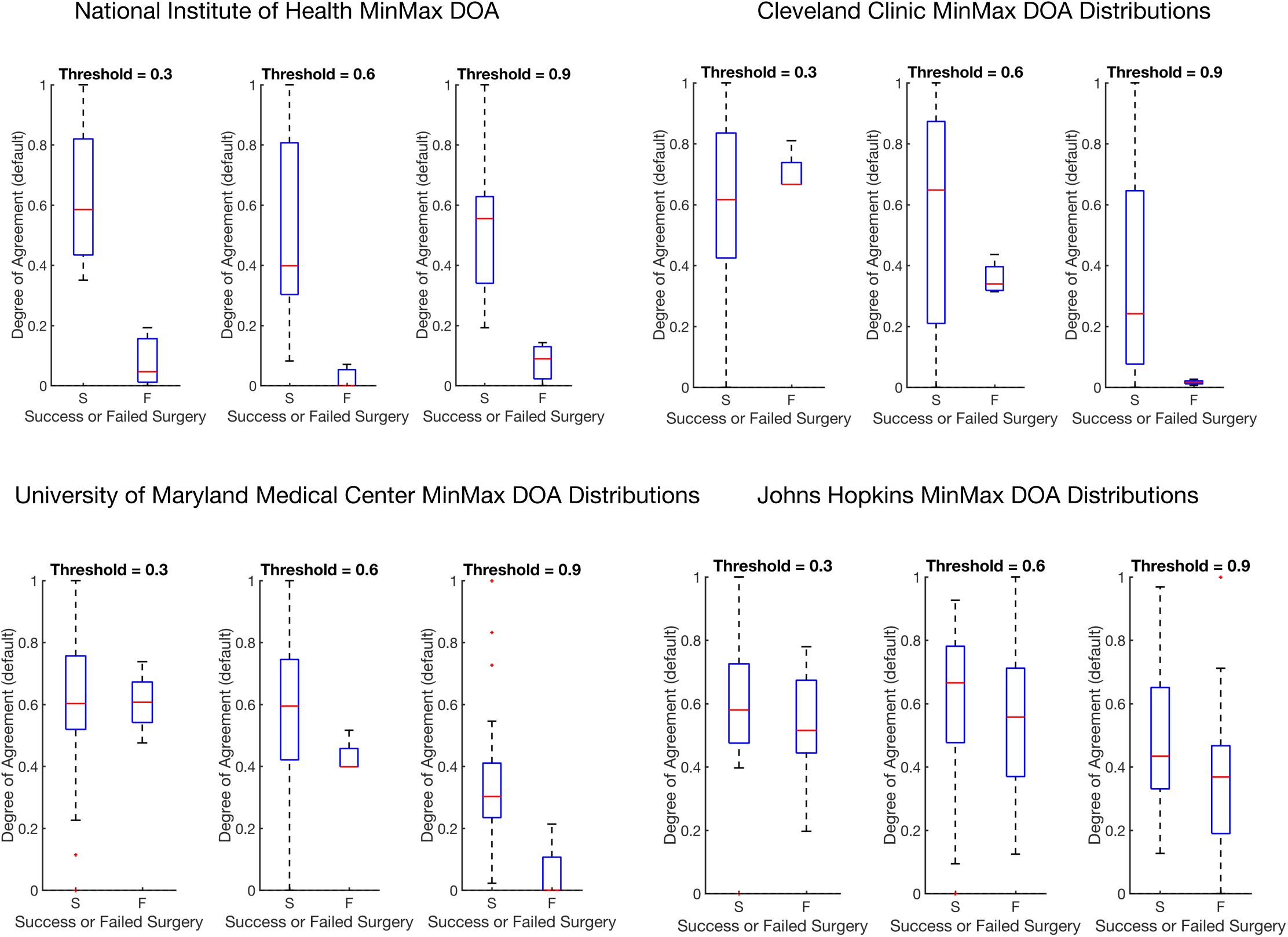
This figure shows distributions of the degree of agreement for every center including JHU after minmax normalization to compare each center on the same scale of success versus failure. Note that minmax normalization scales all distributions between 0 and 1.

In the case that a patient has failed outcomes, we would not expect to see a perfect disagreement DOA score of −1 because of the above reasons. There may have been no visible EZ recorded from the electrode network, or the EZ may not have been fully resected (but part of it was still clinically annotated). It is also important to note that when a patient has a successful surgical outcome, clinicians remove a large portion of the brain, which is a superset of the clinically annotated EZ. It is not certain that all clinically annotated EZ electrodes are actually part of the true underlying EZ, so we would expect some deviation from perfect agreement with the clinically annotated EZ (e.g. we should not expect to see a perfect DOA score of 1 for successful patients).

## Discussion

The definition of the EZ, including its anatomical and electrophysiological signatures, has been an evolving and controversial topic since the foundation of modern epilepsy surgery. The EZ, defined as the site of primary organization of the ictal discharge, refers to the cortical areas connected together through an excessive synchronization at seizure onset (52; 57). Fast activity (FA) at ictal onset has been clinically accepted as the main feature of the EZ since the beginning of invasive monitoring era, particularly in the SEEG literature (52). Since the development of subdural ECoG recordings, much attention has also been paid to the time precedence of phasic transients, especially spiking activities (5; 45). In the last fifteen years, identification of high frequency oscillations (HFO) during interictal and ictal periods in experimental models reoriented research interest towards high-gamma activities in human epilepsies as a potential EZ marker (6; 39; 64). In parallel, DC recordings exemplified the concomitance of ultra-slow and fast frequencies (19; 24; 53; 62).

Although clinical definitions have been explored, a network based operational definition of the EZ is currently not well defined in the literature. Novel computational network analyses may overcome some of the challenges associated with more conventional invasive monitoring recordings methods. In this study, we analyze how centrality signatures of electrode recordings within an epileptic network change over time and how they relate to clinical annotations from four different hospital centers. We take in ECoG and SEEG data from 60 seconds before and after a seizure instance for 42 patients and produce a frequency connectivity network over time using the cross-power spectra of the signal in the 30-90 Hz range. Then we computed the EVC for each electrode at a time window to obtain a normalized ranked centrality of every electrode over time. By overlaying a Gaussian weighting function that was trained only with patients from one center, we then obtain a likelihood for each electrode of being in the EZ. Then we computed a degree of agreement between our algorithm and clinically labeled EZ using the DOA index for all patients by setting an arbitrary threshold.

Some previous approaches for marking the EZ included FA, signal flattening and slow potential shift. Fast activity frequently occurs quasi-simultaneously in multiple areas so that visual discrimination can be cumbersome and lead to subjective interpretations. A different approach, frequency localization, was used by (19). After defining frequencies of interest (FOIs) and plotting their power change over time, they localized the distribution of FOIs in different contacts of the depth electrodes. The EZ, defined as the area exhibiting frequency changes at seizure onset, could then be delineated. In a retrospective and prospective study of patients investigated using SEEG, the same method was applied to test three potential biomarkers of EZ, namely FA, signal flattening and slow potential shift. These biomarkers co-localized with the EZ as defined by standard SEEG criteria and postresection seizure outcome (19).

Other approaches for marking the EZ include HFO analyses. Interictal HFOs have been shown to have some value in identifying the EZ (11; 25; 28; 38; 55; 56). In our comparative analysis, we made modifications to the algorithm based on limitations in the data that was available at the clinical centers. First, in the 1000 Hz sampled data, the number of HFOs is significantly reduced, although the detected HFOs are still useful to identify the EZ (18). The lower sampling rate also required some modifications to the algorithm: the fast-transient artifact detector could not be used (as it requires sampling rates > 2 kHz) and the upper edge on the band-pass filter needed to be reduced from 500 to 400 Hz. Second, the limitation to interictal data restricts the identification of the full EZ: HFO results typically report a very small number of channels involved, which are typically much smaller than the eventual resected volume of tissue. Although HFO analyses show promise in analyzing electrophysiology of epileptic patients, they do not take into account the network nature of epilepsy. HFO analyses are important for analyses of interictal data, since our analysis is limited by requiring recorded seizure events. In future studies, it would be interesting to see how network algorithms and HFO algorithms can complement each other to improve EZ localization.

It is important to note that network-bases analyses is not new to analyzing EEG recordings from epilepsy patients. Previous studies have shown that seizure activity is a dynamic multichannel process and the correlation structure right around a seizure event also follows a typical evolution, similar to our ranked EVC signal (34; 48). In (34; 48), they do not relate it back to EZ, but just look at network dynamics during seizure events. In (47), the authors compute interelectrode synchrony using the mean phase coherence algorithm and relate locally synchronous EEG channels back to the EZ, but only analyzed only 9 patients from a single center. A similar small-scale study was performed in (32) with six epilepsy patients from one center. Other studies use computational models to understand the biophysical mechanisms related to epilepsy surgery (31; 51). In (31), they applied a virtual resection model using data from 10 patients. In (51), the authors developed patient-specific dynamical network models of epileptogenic cortex (computational models). However, there were only 16 patients analyzed from one center.

This manuscript describes a somewhat large-scale research study that applies network-based data analysis tools to invasive EEG data to explore possible EEG signatures of the EZ. In no way are we proposing that this algorithm be directly translated into the clinic. Rather, it now compares how pairwise correlations may improve over quantifying HFOs in each channel individually, which has been the most recently accepted approach. We present a network analysis related back to the annotated EZ, analyzing data from before and after seizures, and analyzing data from multiple centers (with 113 seizures from 42 patients). In our study, we showed that there is a general higher degree of agreement between our algorithm and clinically successful surgical resections of the EZ and a lower degree of agreement between our algorithm and clinically failed resections. By setting a simple threshold on the likelihood maps, we can obtain a similarity measure between our algorithm and clinical labels for both successful and failed surgeries. As the threshold increases, our algorithm becomes better at identifying if successful resections had the correct EZ. We observed that the algorithm’s performance degraded with respect to degree of agreement when patients were implanted sparsely with single strips across all four lobes (UMMC patients) and sometimes in both hemispheres. The clinicians place these strips with such wide coverage if there is no clear pre-implantation hypothesis and if seizures are thought to be starting from multiple brain regions. Often, these patients do not have clear EZ localization and/or do not end up as candidates for surgery. We also found that if the electrographic onset of seizure is not close to the clinically annotated onset of seizure, then the degree of agreement with clinicians is reduced. The electrographic onset is the start of seizure that is seen on the EEG recordings but not manifested in any behavioral changes in the patient. The clinical onset is the time at which the patient exhibits noticeable behavioral changes due to seizure onset (e.g. muscle twitches).

Our results suggest that network data analytics may be a useful tool to assist in localization of the epileptogenic zone, especially when electrode implantation covers the EZ network densely. This is expected, since the threshold on the network’s likelihood is essentially a threshold on the algorithm’s confidence in an electrode being within the EZ set. Future work entails exploring different weighting functions applied over the rank centrality space and possibly merging features from HFO and network algorithms. Besides looking solely at gamma power (30-90 Hz) cross power matrices, the work could expand to encompass more frequency bands that could contain signals of importance in EZ localization. In addition, a more comprehensive study that compares the outcomes between SEEG and ECoG could help understand limitations of the algorithm, and also be of clinical importance in using SEEG versus ECoG. In addition, if we had more patient data from other centers, then it would be interesting to see how a pooled training procedure may improve our results. This work is meant to supplement the growing evidence in literature that epilepsy is a network phenomena and therefore also requires network algorithms to better understand its manifestation.

## SUPPORTIVE INFORMATION

Code is open source at https://github.com/ncsl/eztrack. Since this was a retrospective data study, there is no table of the JHU patients and their clinical operative notes because the data was not available from JHU.

## ACKNOWLEDGMENTS

AL is supported by NIH T32 EB003383 and SVS by Coulter Foundation and the Maryland Innovation Initiative. R. Y. was supported by the Epilepsy Foundation Predoctoral Research Training Fellowship. S.V.S, N.C, and W.S.A. are supported by Maryland Technology Development Corportation (TEDCO) through MII. S.V.S was supported by the US NSF Career Award 1055560 and the Burroughs Wellcome Fund CASI Award 1007274. Data collection work was supported by the Intramural Research Program at NIH. We would also like to thank the reviewers for their valuable advice.

## AUTHOR CONTRIBUTIONS

S.V.S, J.T.G, A.T, R.Y, S.S and J.G-M helped formulate the project. J.G-M., J.T.G., Z.F., J.B., K.Z., S.I., J.H, C.A., J.H., N.C., E.J., W.S.A. all supplied EEG data. A.L., B.C, S.S., R.Y. and S.V.V contributed to the analyses. S.G., W.S., and R.N. helped perform analysis. Finally, A.L., B.C, S.V.S, and J.G-M. wrote the manuscript.

## REFERENCES

Danielle Smith Bassett and Ed Bullmore. Small-World Brain Networks. The Neuroscientist, 12(6):512–523, dec 2006.

Charles E. Begley, Melissa Famulari, John F. Annegers, David R. Lairson, Thomas F. Reynolds, Sharon Coan, Stephanie Dubinsky, Michael E. Newmark, Cynthia Leibson, E. L. So, and Walter A. Rocca. The Cost of Epilepsy in the United States: An Estimate from Population-Based Clinical and Survey Data. Epilepsia, 41(3):342–351, mar 2000.

Anne T. Berg. Identification of Pharmacoresistant Epilepsy. Neurologic Clinics, 27(4):1003–1013, nov 2009.

Anne T. Berg and Molly M. Kelly. Defining intractability: Comparisons among published definitions. Epilepsia, 47(2):431–436, feb 2006.

Kanokwan Boonyapisit, Imad Najm, George Klem, Zhong Ying, Candice Burrier, Eric LaPresto, Dileep Nair, William Bingaman, Richard Prayson, and Hans Lüders. Epileptogenicity of focal malformations due to abnormal cortical development: Direct electrocorticographic-histopathologic correlations. Epilepsia, 44(1):69–76, jan 2003.

Anatol Bragin, Charles L. Wilson, Richard J. Staba, Mark Reddick, Itzhak Fried and Jerome Engel. Interictal high-frequency oscillations (80-500Hz) in the human epileptic brain: Entorhinal cortex. Annals of Neurology, 52(4):407–415, oct 2002.

Urs Braun, Sarah F Muldoon, and Danielle S Bassett. On Human Brain Networks in Health and Disease. In eLS, volume 22, pages 1–9. John Wiley & Sons, Ltd, Chichester, UK, feb 2015.

M. J. Brodie, S. D. Shorvon, R. Canger, P. Halasz, S. Johannessen, P. Thompson, H. G. Wieser, and P. Wolf. Commission on European Affairs: Appropriate standards of epilepsy care across Europe. Epilepsia, 38(11):1245–1250, nov 1997.

Juan C. Bulacio, Lara Jehi, Chong Wong, Jorge Gonzalez-Martinez, Prakash Kotagal, Dileep Nair, Imad Najm, and William Bingaman. Long-term seizure outcome after resective surgery in patients evaluated with intracranial electrodes. Epilepsia, 53(10):1722–1730, oct 2012.

Edward T. Bullmore and Danielle S. Bassett. Brain Graphs: Graphical Models of the Human Brain Connectome. Annual Review of Clinical Psychology, 7(1):113–140, apr 2011.

Sergey Burnos, Birgit Frauscher, Rina Zelmann, Claire Haegelen, Johannes Sarnthein, and Jean Gotman. The morphology of high frequency oscillations (HFO) does not improve delineating the epileptogenic zone. Clinical Neurophysiology, 127(4):2140–2148, apr 2016.

Samuel P. Burns, Sabato Santaniello, Robert B. Yaffe, Christophe C. Jouny, Nathan E. Crone, Gregory K. Bergey, William S. Anderson, and Sridevi V. Sarma. Network dynamics of the brain and influence of the epileptic seizure onset zone. Proceedings of the National Academy of Sciences, 111(49):E5321–E5330, dec 2014.

Lorena Deuker, Edward T. Bullmore, Marie Smith, Soren Christensen, Pradeep J. Nathan, Brigitte Rockstroh, and Danielle S. Bassett. Reproducibility of graph metrics of human brain functional networks. NeuroImage, 47(4):1460–1468, oct 2009.

Mark A. Ferro and Kathy N. Speechley. Depressive symptoms among mothers of children with epilepsy: A review of prevalence, associated factors, and impact on children. Epilepsia, 50(11):2344–2354, nov 2009.

Robert S. Fisher. Therapeutic devices for epilepsy. Annals of Neurology, 71(2):157–168, feb 2012.

F. Gilliam, R. Kuzniecky, K. Meador, R. Martin, S. Sawrie, M. Viikinsalo, R. Morawetz, and E. Faught. Patient-oriented outcome assessment after temporal lobectomy for refractory epilepsy. Neurology, 53(4):687–687, sep 1999.

F G Gilliam. Diagnosis and treatment of mood disorders in persons with epilepsy, volume 18. Rapid Science Publishers, 2005.

Stephen V. Gliske, Zachary T. Irwin, Kathryn A. Davis, Kinshuk Sahaya, Cynthia Chestek, and William C. Stacey. Universal automated high frequency oscillation detector for real-time, long term EEG. Clinical Neurophysiology, 127(2):1057–1066, 2016.

Vadym Gnatkovsky, Marco De Curtis, Chiara Pastori, Francesco Cardinale, Giorgio Lo Russo, Roberto Mai, Lino Nobili, Ivana Sartori, Laura Tassi, and Stefano Francione. Biomarkers of epileptogenic zone defined by quantified stereo-EEG analysis. Epilepsia, 55(2):296–305, feb 2014.

Jorge Gonzalez-Martinez, Juan Bulacio, Andreas Alexopoulos, Lara Jehi, William Bingaman, and Imad Najm. Stereoelectroencephalography in the “difficult to localize” refractory focal epilepsy: Early experience from a North American epilepsy center. Epilepsia, 54(2):323–330, feb 2013.

Jorge Gonzalez´Mart´ınez, Juan Bulacio, Susan Thompson, John Gale, Saksith Smithason, Imad Najm, and William Bingaman. Technique, results, and complications related to robot-assisted stereoelectroencephalography. Neurosurgery, 78(2):169–179, 2016.

Jean Gotman. Measurement of small time differences between EEG channels: Method and application to epileptic seizure propagation. Electroencephalography and Clinical Neurophysiology, 56(5):501–514, 1983.

Bruce P. Hermann, Michael Seidenberg, Christian Dow, Jana Jones, Paul Rutecki, Abhik Bhattacharya, and Brian Bell. Cognitive prognosis in chronic temporal lobe epilepsy. Annals of Neurology, 60(1):80–87, jul 2006.

Akio Ikeda, Kiyohito Terada, Nobuhiro Mikuni, Richard C. Burgess, Youssef Comair, Waro Taki, Toshiaki Hamano, Jun Kimura, Hans O. Lüders, and Hiroshi Shibasaki. Subdural recording of ictal DC shifts in neocortical seizures in humans. Epilepsia, 37(7):662–674, jul 1996.

J. Jacobs, R. Staba, E. Asano, H. Otsubo, J. Y. Wu, M. Zijlmans, I. Mohamed, P. Kahane, F. Dubeau, V. Navarro, and J. Gotman. High-frequency oscillations (HFOs) in clinical epilepsy. Progress in Neurobiology, 98(3):302–315, sep 2012.

L. E. Jeha, I. M. Najm, W. E. Bingaman, F. Khandwala, P. Widdess-Walsh, H. H. Morris, D. S. Dinner, D. Nair, N. Foldvary-Schaeffer, R. A. Prayson, Y. Comair, R. O’Brien, J. Bulacio, A. Gupta, and H. O. Lüders. Predictors of outcome after temporal lobectomy for the treatment of intractable epilepsy. Neurology, 66(12):1938–1940, jun 2006.

Lara E. Jeha, Imad Najm, William Bingaman, Dudley Dinner, Peter Widdess-Walsh, and Hans Lüders. Surgical outcome and prognostic factors of frontal lobe epilepsy surgery. Brain, 130(2):574–584, feb 2007.

Jing Xiang. Localizing Functional Brain Cortices and Epileptogenic Zones With HFOs. Technical report, Children’s Hospital Medical Center, Cincinnati, 2008.

Won Young Jung, Steven V. Pacia, and Orrin Devinsky. Neocortical Temporal Lobe Epilepsy: Intracranial EEG Features and Surgical Outcome, 1999.

Matthew S D Kerr, Samuel P. Burns, John Gale, Jorge Gonzalez-Martinez, Juan Bulacio, and Sridevi V. Sarma. Multivariate analysis of SEEG signals during seizure. Proceedings of the Annual International Conference of the IEEE Engineering in Medicine and Biology Society, EMBS, pages 8279–8282, 2011.

Ankit N. Khambhati, Kathryn A. Davis, Timothy H. Lucas, Brian Litt, and Danielle S. Bassett. Virtual Cortical Resection Reveals Push-Pull Network Control Preceding Seizure Evolution. Neuron, 91(5):1170–1182, 2016.

A. Korzeniewska, M. C. Cervenka, C. C. Jouny, J. R. Perilla, J. Harezlak, G. K. Bergey, P. J. Franaszczuk, and N. E. Crone. Ictal propagation of high frequency activity is recapitulated in interictal recordings: Effective connectivity of epileptogenic networks recorded with intracranial EEG. NeuroImage, 101:96–113, 2014.

M. A. Kramer, U. T. Eden, E. D. Kolaczyk, R. Zepeda, E. N. Eskandar, and S. S. Cash. Coalescence and Fragmentation of Cortical Networks during Focal Seizures. Journal of Neuroscience, 30(30):10076–10085, 2010.

Mark A. Kramer, Eric D. Kolaczyk, and Heidi E. Kirsch. Emergent network topology at seizure onset in humans. Epilepsy Research, 79(2–3):173–186, 2008.

Patrick Kwan and Martin J. Brodie. Early Identification of Refractory Epilepsy. New England Journal of Medicine, 342(5):314–319, feb 2000.

Hans O. Lüders, Imad Najm, Dileep Nair, Peter Widdess-Walsh, and William Bingman. The epileptogenic zone: General principles. Epileptic Disorders, 8(SUPPL. 2), 2006.

K. A. Ludwig, R. M. Miriani, N. B. Langhals, M. D. Joseph, D. J. Anderson, and D. R. Kipke. Using a Common Average Reference to Improve Cortical Neuron Recordings From Microelectrode Arrays. Journal of Neurophysiology, 101(3):1679–1689, mar 2009.

Urszula Malinowska, Gregory K. Bergey, Jaroslaw Harezlak, and Christophe C. Jouny. Identification of seizure onset zone and preictal state based on characteristics of high frequency oscillations. Clinical Neurophysiology, 126(8):1505–1513, aug 2015.

A. Matsumoto, B. H. Brinkmann, S. Matthew Stead, J. Matsumoto, M. T. Kucewicz, W. R. Marsh, F. Meyer, and G. Worrell. Pathological and physiological high-frequency oscillations in focal human epilepsy. Journal of Neurophysiology, 110(8):1958–1964, 2013.

Anne M. McIntosh, Renate M. Kalnins, L. Anne Mitchell, Gavin C A Fabinyi, Regula S. Briellmann, and Samuel F. Berkovic. Temporal lobectomy: Long-term seizure outcome, late recurrence and risks for seizure recurrence. Brain, 127(9):2018–2030, aug 2004.

Miranda I. Murray, Michael T. Halpern, and Ilo E. Leppik. Cost of refractory epilepsy in adults in the USA. Epilepsy Research, 23(2):139–148, 1996.

Dileep R. Nair, Richard Burgess, Cameron C. McIntyre, and Hans Lüders. Chronic subdural electrodes in the management of epilepsy. Clinical Neurophysiology, 119(1):11–28, 2008.

Ernst Niedermeyer and F H Lopes Da Silva. Electroencephalography: Basic Principles, Clinical Applications, and Related Fields, volume 1. Urban & Schwarzenberg, 2004.

C¸agatay Önal, Hiroshi Otsubo, Takashi Araki, Shiro Chitoku, Ayako Ochi, Shelly Weiss, William Logan, Irene Elliott, O. Carter Snead, and James T. Rutka. Complications of invasive subdural grid monitoring in children with epilepsy. Journal of Neurosurgery, 98(5):1017–1026, may 2003.

Andreé Palmini, Antonio Gambardella, Frederick Andermann, Francois Dubeau, Jaderson C. da Costa, Andreé Olivier, Donatella Tampieri, Pierre Gloor, Felipe Quesney, Eva Andermann, Eduardo Paglioli, Eliseu PaglioliNeto, Ligia Coutinho Andermann, Richard Leblanc, and HyoungIhl I Kim. Intrinsic epileptogenicity of human dysplastic cortex as suggested by corticography and surgical results. Annals of Neurology, 37(4):476–487, apr 1995.

Sabato Santaniello, Samuel P. Burns, Alexandra J. Golby, Jedediah M. Singer, William S. Anderson, and Sridevi V. Sarma. Quickest detection of drug-resistant seizures: An optimal control approach. Epilepsy and Behavior, 22(SUPPL. 1), 2011.

C. A. Schevon, J. Cappell, R. Emerson, J. Isler, P. Grieve, R. Goodman, G. Mckhann, H. Weiner, W. Doyle, R. Kuzniecky, O. Devinsky, and F. Gilliam. Cortical abnormalities in epilepsy revealed by local EEG synchrony. NeuroImage, 35(1):140–148, 2007.

Kaspar Schindler, Howan Leung, Christian E. Elger, and Klaus Lehnertz. Assessing seizure dynamics by analysing the correlation structure of multichannel intracranial EEG. Brain, 130(1):65–77, 2007.

Stephan U. Schuele and Hans O. Lüders. Intractable epilepsy: management and therapeutic alternatives. The Lancet Neurology, 7(6):514–524, 2008.

Siew Ju See, Lara E. Jehi, Sumeet Vadera, Juan Bulacio, Imad Najm, and William Bingaman. Surgical outcomes in patients with extratemporal epilepsy and subtle or normal magnetic resonance imaging findings. Neurosurgery, 73(1):68–76, jul 2013.

Nishant Sinha, Justin Dauwels, Marcus Kaiser, Sydney S. Cash, M. Brandon Westover, Yujiang Wang, and Peter N. Taylor. Predicting neurosurgical outcomes in focal epilepsy patients using computational modelling. Brain, 140(2):319–332, 2017.

J. Talairach and J. Bancaud. Stereotaxic Approach to Epilepsy. In Progress in neurological surgery, volume 5, pages 297–354. Karger Publishers, 1973.

P. J. Thompson, H. Conn, S. A. Baxendale, E. Donnachie, K. McGrath, C. Geraldi, and J. S. Duncan. Optimizing memory function in temporal lobe epilepsy. Seizure, 38:68–74, may 2016.

Horst Urbach, Jörg Hattingen, Joachim Von Oertzen, Cordelia Luyken, Hans Clusmann, Thomas Kral, Martin Kurthen, Johannes Schramm, Ingmar Blümcke, and Hans H. Schild. MR imaging in the presurgical workup of patients with drug-resistant epilepsy. American Journal of Neuroradiology, 25(6):919–926, 2004.

Naotaka Usui, Kiyohito Terada, Koichi Baba, Kazumi Matsuda, Fumihiro Nakamura, Keiko Usui, Miyako Yamaguchi, Takayasu Tottori, Shuichi Umeoka, Shigeru Fujitani, Akihiko Kondo, Tadahiro Mihara, and Yushi Inoue. Clinical significance of ictal high frequency oscillations in medial temporal lobe epilepsy. Clinical Neurophysiology, 122(9):1693–1700, sep 2011.

N. E C Van Klink, M. A. Van’t Klooster, R. Zelmann, F. S S Leijten, C. H. Ferrier, K. P J Braun, P. C. Van Rijen, M. J A M Van Putten, G. J M Huiskamp, and M. Zijlmans. High frequency oscillations in intra-operative electrocorticography before and after epilepsy surgery. Clinical Neurophysiology, 125(11):2212–2219, nov 2014.

Fabrice Wendling, Patrick Chauvel, Arnaud Biraben, and Fabrice Bartolomei. From Intracerebral EEG Signals to Brain Connectivity: Identification of Epileptogenic Networks in Partial Epilepsy. Frontiers in Systems Neuroscience, 4, 2010.

Elise Whitley and Jonathan Ball. The sign test. Critical care (London, England), 6(6):509–13, dec 2002.

P. Widdess-Walsh, L. Jeha, D. Nair, P. Kotagal, W. Bingaman, and I. Najm. Subdural electrode analysis in focal cortical dysplasia: Predictors of surgical outcome. Neurology, 69(7):660–667, aug 2007.

Heinz Gregor Wieser. Epilepsy surgery: Past, present and future. Seizure, 7(3):173–184, jun 1998.

Greg A. Worrell, Landi Parish, Stephen D. Cranstoun, Rachel Jonas, Gordon Baltuch, and Brian Litt. High-frequency oscillations and seizure generation in neocortical epilepsy. Brain, 127(7):1496–1506, jul 2004.

L. Wu and J. Gotman. Segmentation and classification of EEG during epileptic seizures. Electroencephalography and Clinical Neurophysiology, 106(4):344–356, 1998.

Robert Yaffe, Sam Burns, John Gale, Hyun Joo Park, Juan Bulacio, Jorge Gonzalez-Martinez, and Sridevi V. Sarma. Brain state evolution during seizure and under anesthesia: A network-based analysis of stereotaxic eeg activity in drug-resistant epilepsy patients. In Proceedings of the Annual International Conference of the IEEE Engineering in Medicine and Biology Society, EMBS, pages 5158–5161. IEEE, aug 2012.

Maeike Zijlmans, Premysl Jiruska, Rina Zelmann, Frans S S Leijten, John G R Jefferys, and Jean Gotman. High-frequency oscillations as a new biomarker in epilepsy. Annals of Neurology, 71(2):169–178, feb 2012.

